# Multiple Receptor Tyrosine Kinases Regulate Dengue Infection of Hepatocytes

**DOI:** 10.1101/2023.07.30.549949

**Authors:** Natasha M. Bourgeois, Ling Wei, Nhi N. T. Ho, Maxwell L. Neal, Denali Seferos, Tinotenda Tongogara, Fred D. Mast, John D. Aitchison, Alexis Kaushansky

**Affiliations:** Department of Global Health, University of Washington, Seattle WA 98195, USA; Center for Global Infectious Disease Research, Seattle Children’s Research Institute, Seattle WA 98109, USA; Department of Pediatrics, University of Washington, Seattle WA 98195, USA

**Author notes:** Nexelis Pharmaceuticals, Seattle WA 98199, USA.

## Abstract

Dengue is an arboviral disease causing severe illness in over 500,000 people each year. Currently, there is no way to constrain dengue in the clinic. Host kinase regulators of dengue virus (DENV) infection have the potential to be disrupted by existing therapeutics to prevent infection and/or disease progression. To evaluate kinase regulation of DENV infection, we performed kinase regression (KiR), a machine learning approach that predicts kinase regulators of infection using existing drug-target information and a small drug screen. We infected hepatocytes with DENV *in vitro* in the presence of a panel of 38 kinase inhibitors then quantified the effect of each inhibitor on infection rate. We employed elastic net regularization on these data to obtain predictions of which of 300 kinases are regulating DENV infection. Thirty-six kinases were predicted to have a functional role. Intriguingly, seven of the predicted kinases – EPH receptor A4 (EPHA4), EPH receptor B3 (EPHB3), EPH receptor B4 (EPHB4), erb-b2 receptor tyrosine kinase 2 (ERBB2), fibroblast growth factor receptor 2 (FGFR2), Insulin like growth factor 1 receptor (IGF1R), and ret proto-oncogene (RET) – belong to the receptor tyrosine kinase (RTK) family, which are already therapeutic targets in the clinic. We demonstrate that predicted RTKs are expressed at higher levels in DENV infected cells. Knockdown of ERBB2, FGFR2 and IGF1R reduces DENV infection in hepatocytes. Finally, we observe differential temporal induction of ERBB2 and IGF1R following DENV infection, highlighting their unique roles in regulating DENV. Collectively, our findings underscore the significance of multiple RTKs in DENV infection and advocate further exploration of RTK-oriented interventions against dengue.

## INTRODUCTION

Dengue is a neglected tropical disease caused by dengue virus (DENV), a mosquito-borne flavivirus [1]. Dengue incidence has increased at an alarming rate, with over 4.2 million dengue cases reported to the World Health Organization in 2019 compared to the 500,000 cases reported in the year 2000 [2]. Strikingly, it is estimated that hundreds of millions more dengue cases are evading surveillance each year [3].

Global warming and urbanization are expanding suitable habitats for mosquito vector populations, lending to the prediction of further escalated dengue incidence in the coming years [4]. In the absence of an effective vaccine or specific therapeutics for DENV, instances of severe disease have also been rising [2]. Identifying effective interventions against infection is imperative to combat the growing global health burden of dengue.

Efforts towards identifying compounds that directly target DENV proteins are ongoing. DENV comprises an enveloped positive-sense single-stranded RNA genome which encodes three structural proteins – envelope (Env), pre-membrane (PrM), and capsid (C) – and seven non-structural proteins – NS1, NS2A, NS2B, NS3, NS4A, NS4B, and NS5 (reviewed in [5]). Many compounds targeting these proteins exhibit efficacy *in vitro* and *in vivo* (reviewed in [6]). However, none of these have demonstrated efficacy against viral load or disease in clinical trials [7; 8; 9; 10].

On the other hand, host-targeting compounds have provided some protection against dengue disease in recent clinical trials [11]. Importantly, targeting the host has proven to be a successful therapeutic strategy for other infectious diseases, including human papilloma virus, hepatitis C virus, hepatitis B virus, and human immunodeficiency virus [12; 13]. In addition to curbing disease in clinical trials, this strategy has also demonstrated promise for blocking DENV infection, with synergistic effects demonstrated against viral load *in vitro* and *in vivo* when combining a host-targeting α-glucosidase inhibitor and the broad antiviral ribavirin, a guanosine analog [14]. However, a comprehensive understanding of druggable host molecules that are critical for successful DENV infection is unavailable, limiting the pool of candidates for host targeted intervention against dengue.

Ample evidence demonstrates that DENV relies on host factors for infection and pathogenesis, as extensively reviewed in [15]. For instance, DENV requires host attachment receptors and regulators of endocytosis for entry [16; 17; 18; 19; 20; 21], exploits regulators and structural components of host translation machinery for viral genome replication [22; 23; 24; 25], and manipulates factors involved in the immune response [26; 27; 28; 29]. Notably, protein kinases serve as upstream regulators of each of these events. Taken together with the existence of hundreds of kinase-targeting drugs already existing in the clinic [30], kinases are promising candidate targets for DENV therapeutics.

Evidence for a role of kinase activity in DENV infection and pathogenesis continues to accumulate [31; 32; 33; 34; 35], and multiple kinase inhibitors have been shown to restrict dengue infection *in vitro* and *in vivo* [36; 37; 38; 39; 40]. Despite numerous high-throughput screens aiming to identify host factors regulating DENV [31; 41; 42; 43; 44; 45; 46], these attempts have failed to identify kinase regulators of infection, perhaps due to the extensive compensatory roles of other kinases or insufficient degree of depletion. Here, we overcome this shortcoming of prior screening methods by employing Kinase Regression (KiR) on DENV infection. KiR is an innovative computational tool that uses a panel of 38 kinase inhibitors – with known enzymatic inhibition activity against a total of 300 kinases spanning the human kinome – as chemical probes to screen kinase activity regulating infection [47; 48]. KiR uses elastic net regularization to decipher which shared targets of the inhibitor panel influence infection (Figure 1A). Notably, this approach has been successfully utilized to identify key host factors regulating malaria liver infection and disruption of the blood-brain barrier [49; 50].

**Figure 1.**
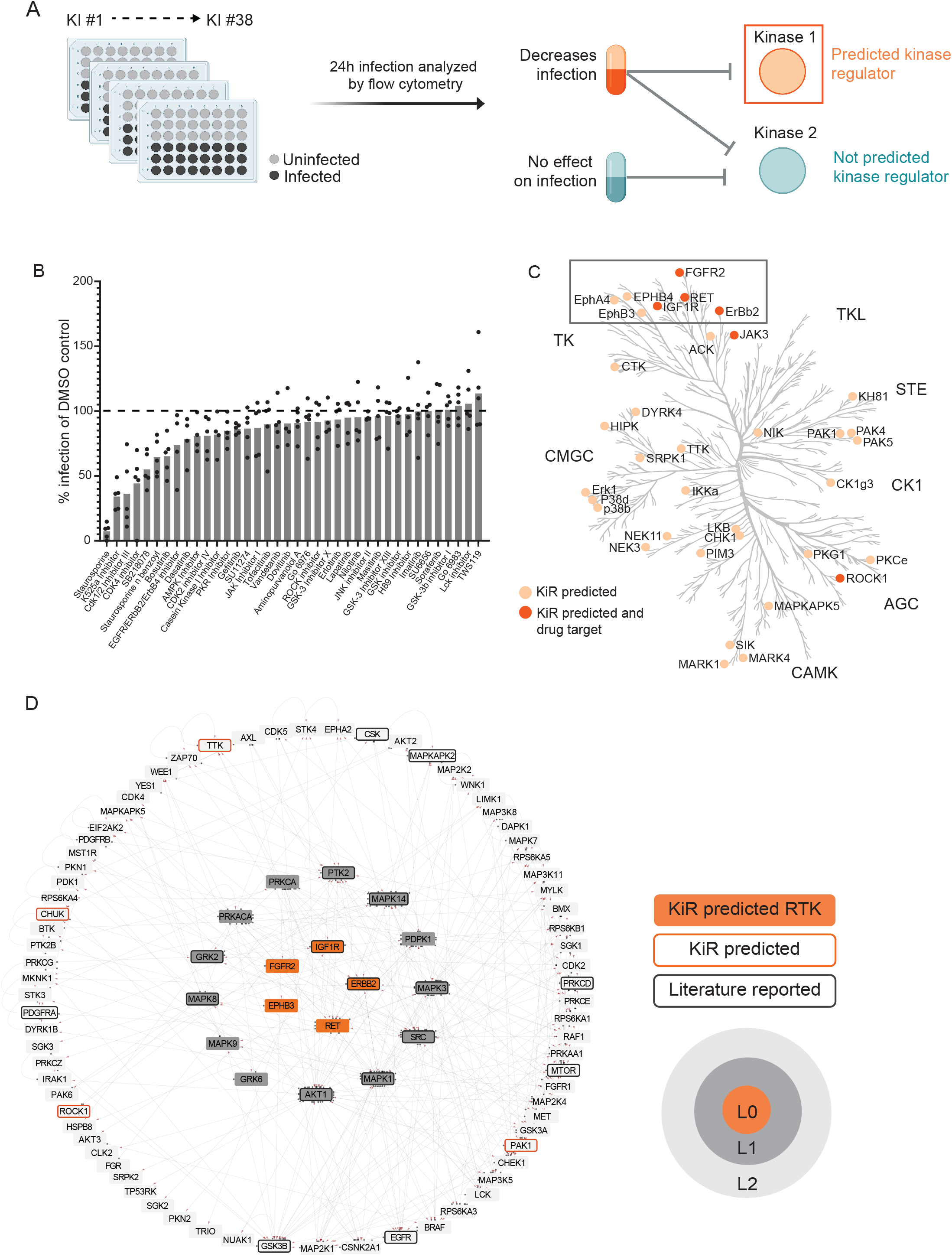
Kinase Regression (KiR) on dengue (DENV) infection predicts seven receptor tyrosine kinases (RTKs) as regulators of infection. (A) HepG2 cells were pre-treated with KiR kinase inhibitor (KI) panel (Supplementary Table 1) at a concentration of 500 nM for one hour, followed by infection with DENV2 MON601 at a MOI of 2 under continuous inhibitor exposure. 24 h post-infection, cells were fixed and stained with antibody against DENV envelope (Env) protein. Flow cytometry analysis subsequently determined the percentage of cells staining positively for Env. (B) Percent difference in infection rates in response to each inhibitor compared to the mock-treated control (DMSO). Data below the dashed horizontal line indicates lower infection rates than the mock control and vice versa. (C) KiR predicted 36 kinases as potential regulators of dengue infection in HepG2s. The phylogenetic kinase family tree depicts KiR-predicted kinases and the subset that are targets of existing therapeutics in light or dark orange, respectively. (D) Interaction network denotes predicted RTKs (in orange), their the direct (L1, gray) or indirect (L2, light gray) interactors, other predicted kinases (orange outline), and kinases implicated in existing DENV literature (gray outline). The directionality of interactions is shown, where gray circles indicate source kinase and red arrows indicate substrate.

We performed KiR on DENV infection in HepG2 cells and predicted 36 kinase regulators. We further investigated a subset of these predictions, namely the receptor tyrosine kinases (RTKs), since many predicted kinases in this family are targeted by drugs that are already in clinical use. To validate and characterize their involvement during DENV, we determined the impact of DENV infection on their expression and activity, and we tested the impact of their knockdown on multiple DENV factors. We show that levels of KiR-predicted kinases are significantly higher in infected cells both in total and at the surface. Knockdown of a subset of these kinases significantly disrupts DENV infection, and their phosphorylation is induced at different times throughout infection. Our findings provide support for further investigation into targeting RTKs as a as a promising avenue for combating dengue infection and pathogenesis.

## MATERIALS AND METHODS

### Cell Culture and Maintenance

All cells used in this study were propagated from commercial passage 0 stock and stored at low passage in 10-90% FBS/10% DMSO in LN_2_. Sterility of cell cultures was maintained through the use of a biosafety cabinet. Cell cultures were validated for absence of mycoplasma contamination before, during, and after experimental work using the MycoStrip™ - Mycoplasma Detection Kit (Invivogen #rep-mys). All experimental cell lines were grown under specific conditions and handled following precise procedures to ensure optimal growth and reproducibility. Typing and authentication for each cell line was provided by the American Type Culture Collection (ATCC). A maximum of 25 passages was maintained for each cell line.

HepG2 cells were received from ATCC (#HB-8065) and grown in primocin (100 µg/mL)-supplemented Complete Hepatocyte Media (CHM): DMEM Glutamax (Gibco™ #10566016) with 10% heat-inactivated FBS (SeraPrime #F31016HI), and 4 mM L-glutamine filtered through a 0.22 µM polyethersulfone (PES) membrane. Cells were maintained in tissue culture-treated flasks at 37 °C, 5% CO_2_. Upon reaching 90% confluency, cells were washed once with 1X PBS then detached with 0.25% Trypsin-EDTA (Gibco™ #25200072) for 5 min at 37 °C, 5% CO_2_. Detached cells were centrifuged at 158 rcf for 3 min to remove trypsin, resuspended in primocin-supplemented CHM, then filtered through a 40 µM nylon mesh cell strainer to reduce cell clumps for improved growth and counting consistency.

Vero cells were received from ATCC (#CCL-81) and grown in primocin (100 µg/mL)-supplemented Complete Vero Media (CVM): DMEM Glutamax with 10% FBS, 1X MEM Non-Essential Amino Acid solution (NEAA, Gibco™ #11140050) and filtered through a 0.22 µM PES membrane. Cells were grown in tissue culture-treated flasks at 37 °C, 5% CO_2_. Upon reaching 100% confluency, cells were washed once with 1X PBS then detached with 0.25% Trypsin-EDTA for 5 min at 37 °C, 5% CO_2_. Detached cells were centrifuged at 158 rcm for 3 min to remove trypsin then resuspended in primocin-supplemented CVM.

C6/36 cells were received from ATCC (#CRL-1660) and grown in Complete C6/36 Media (CCM): MEM with 10% FBS, 1X MEM NEAA, and 100 µg/mL primocin filtered through a 0.22 µM PES membrane. Cells were grown in tissue culture-treated flasks at 28 °C, 5% CO_2_. Upon reaching 100% confluency, cells were washed once with 1X PBS then detached with 0.25% Trypsin-EDTA for 5 min at 28 °C, 5% CO_2_. Detached cells were centrifuged at 158 rcf for 3 min to remove trypsin then resuspended in CCM HEK293FT cells were a gift from Alan Aderem (Seattle Children’s Research Institute) and grown in primocin (100 µg/mL)-supplemented Complete HEK Media (CHKM): DMEM with 10% FBS, 25 mM HEPES, 1X MEM NEAA and filtered through a 0.22 µM PES membrane. Cells were grown in tissue culture-treated flasks at 37 °C, 5% CO_2_. Upon reaching 100% confluency, cells were gently washed once with 1X PBS then detached with 0.025% Trypsin-EDTA (diluted in 1X PBS) for 5 min at room temperature. Detached cells were centrifuged at 158 rcf for 3 min to remove trypsin then resuspended in primocin-supplemented CHKM.

### shRNA-mediated Gene Knockdown

Non-replicating shRNA lentivirus were generated in HEK293FT cells transfected with MISSION plasmids procured from Sigma-Aldrich (details in Supplementary Table 2). Transfection involved the combination of MISSION plasmid (6 µg), pCMV-VSV-G (3 mg/ml), psPax2 (6 mg/ml), and Polyethylenimine Hydrochloride (PEI Max, 1 mg/mL) in 500 µl serum free-DMEM. The resulting solution was vortexed and incubated at room temperature for 10 min before being added dropwise to HEK293FT cells at 70% confluency in 10 cm^2^ TC-treated dishes. Following overnight incubation at 37 °C, 5% CO_2_, media was replaced, and cells were incubated overnight. Lentivirus-containing supernatant was harvested and filtered through a 0.45 µm PVFD membrane over the subsequent two days.

For the knock-down of host kinases of interest, HepG2 cells were reverse-transduced with the generated shRNA lentivirus. Detached HepG2 cells were mixed with lentivirus (1 ml lentivirus-containing supernatant per 4×10^6^ cells) in antibiotic-free CHM supplemented with 1 µg/ml polybrene (EMD Millipore TR-1003-G) then plated in 10 cm^2^ TC-treated dishes. Following overnight incubation at 37 °C, 5% CO2, cells were replenished with CHM then incubated overnight. For the next seven days, transduced cells were selected in CHM supplemented with 1 µg/ml puromycin (replenished daily). Non-transduced cells were always included in parallel as a positive control for puromycin killing. After puromycin selection, knockdown was verified at the protein level and cells were utilized for experimentation.

### Viral Production, Propagation, and Storage

DENV2 MON601, a molecular clone of DENV-2 New Guinea strain C, was generated by transfecting *in vitro*-transcribed RNA into Vero cells [51]. Virus was propagated by interchangeably infecting 80% confluent C6/36 or Vero cell monolayers with low-passage stock virus at an MOI of 0.01 in Dengue Stock Media (DSM), comprising DMEM Glutamax supplemented with 2% FBS, 1X MEM NEAA, 25 mM HEPES, and 4 mM L-glutamine. Supernatants, collected twice weekly between 5-14 days post-infection, were cleared of cellular debris by centrifugation at 632 rcf followed by filtration through a 0.2 µM CA filter and stored at −80 °C. The viral titer of stocks was enumerated as described below.

### Viral Stock Titer Quantification

Virus stocks were titrated using a flow cytometry approach as previously described [52]. Briefly, Vero cells were infected with serially diluted virus stocks and analyzed at 24 hours post-infection by flow cytometry. Fixed cells were stained with antibody specific to flavivirus envelope protein (4G2) conjugated to AlexaFluor488 and the percentage of cells that stained positive for 4G2-AlexaFluor488 was quantified. The titer, expressed as fluorescence forming units (FFU) per m of virus stock, was derived from the percentage of infected cells relative to the virus volume and cell count. The titer was used to calculate the volume of virus stock necessary to achieve desired MOI.

### Viral Infection

For experimental infections, plated cells were washed once with 1X PBS then titered virus stock diluted in Opti-MEM to an MOI of 2 was added to cells at the minimum volume per well. Cells were incubated for 90 min at 37 °C, 5% CO_2_ then washed once with 1X PBS. Cells were then replenished with DSM and incubated at 37°C, 5% CO_2_ for the indicated time (initial addition of virus is the start time).

### DENV Detection

Pan-flavivirus envelope (Env) antibodies were prepared from hybridoma 290 supernatants and purified by protein A/G chromatography [32]. Anti-Env was conjugated to Alexa Fluor 488 (Env-488) and titrated to determine the optimal concentration between 1:500 and 1:5000. Antibodies against DENV non-structural protein 3 (NS3) were obtained from GeneTex (GTX124252), conjugated to Alexa Fluor 647 (NS3-647), and titrated as with 4G2-488. Fluorophores were conjugated to these primary antibodies using Thermo Scientific™ Antibody Labeling Kits according to the manufacturers protocol.

### RTK Detection

Primary antibodies against EphA4 (Thermo Scientific™ #PA5-14578), EphB3 (Santa Cruz Biotechnology, Inc. #sc-100299), EphB4 (GeneTex #GTX108595), ErbB2 (Cell Signaling Technology, Inc. (CST) # 2165), FGFR2 (CST #11835), IGF-1R (R&D Systems #MAB391) and RET (CST #3220) were used at a concentration of 1:100 then stained with anti-rabbit-PE or anti-mouse Pacific Blue secondary antibody (BioLegend® #406421, Life Technologies #P10993) for flow cytometry.

For western blots, primary RTK antibodies were used at a 1:1000 concentration followed by anti-rabbit or -mouse-HRP (R&D Systems #HAF008, #HAF007). Primary antibodies p-ErbB2 (MilliporeSigma #04-293) and p-IGF-1R (MilliporeSigma #ABE332) were used at a 1:1000 concentration, p-FGFR2 (CST #3471) was used at a 1:500 concentration. Anti-mouse or anti-rabbit GAPDH were used for loading control staining.

### Flow Cytometry

To harvest cells for flow cytometry, HepG2s were washed once with 1X PBS then treated with TrypLE (Gibco™ #12604021) and incubated for 5 min at 37 °C, 5% CO_2_. Detached cells were diluted in CHM, transferred to a 96-well U-bottom plate, then centrifuged at 158 rcf for 5 min to pellet. The supernatant was discarded and the pellet was resuspended in 3.7% paraformaldehyde (PFA, VWR #100503-917) then incubated at room temperature on a shaker for 15 min for chemical fixation. Fixed cells were further prepared and assayed for flow cytometry as described below or stored at 4 °C for no more than one week.

For analysis of total protein levels in cells, fixed cells were washed twice with 1X PBS then incubated in 0.1% Triton-X-100/1X PBS at room temperature on a shaker for 10 min. Permeabilized cells were washed twice with 1X PBS then resuspended in 0.01% Triton-X-100/2% BSA/1X PBS (GoldBio #A-420) at room temperature on a shaker for 1 hour or overnight at 4 °C to block non-specific protein binding. Blocked cells were pelleted at 632 rcf for 5 min and then resuspended in indicated detection antibody. Cells were stained for at least 2 h at room temperature or overnight at 4 °C. For cells stained with unconjugated primary, cells were washed twice with 1X PBS after primary antibody incubation and then resuspended in secondary antibody solution for 1-2 h at room temperature. For analysis of surface proteins levels in cells, fixed cells were washed twice with 1X PBS then resuspended in 2% BSA/1X PBS to block non-specific binding. After staining for kinase detection as described above, cells were permeabilized, blocked, and stained for DENV. Stained cells were washed twice with 1X PBS then assayed on an 18-color FACS analyzer harboring 405, 488, 532 and 640 nm lasers.

The collected events were analyzed using FlowJo. Events were gated for cells by size on FSC-A x SSC-A (Supplementary Figure 1A). Cells were gated for infection using uninfected sample stained with anti-Env-488 or anti-NS3-647 (Supplementary Figure 1B-C). The percentage of cells stained positive for Env-488 or NS3-647 is denoted as % infection. Kinase protein levels were obtained by adding the Geometric Mean statistic to the relevant fluorescence channel and denoted as Mean Fluorescence Intensity (MFI).

### Western Blot

To collect cells for analysis by western blot, HepG2s were washed once with 1X PBS then lysed in SDS lysis buffer (50 mM Tris HCl/2% SDS/5% glycerol/5 mM EDTA/1mM NaF/dH_2_O) supplemented with cOmplete Protease Inhibitor Cocktail Tablets (Roche #11836170001), Phosphatase Inhibitor Cocktail 2 (Sigma-Aldrich® #P5726), B-GP (Sigma-Aldrich® #G9422), PMSF (Sigma-Aldrich® #10837091001), Na_3_VO_4_ (Sigma-Aldrich® #567540-5GM) and DTT (Sigma-Aldrich® # D9779). Lysates were transferred to QIAshredder tubes (QIAGEN #79656) then centrifuged at 1935 rcf for 5 min to separate proteins from genetic material. Bolt Sample Reducing Agent (Invitrogen™ B0009) was added to the eluate and incubated at 70 °C for 10-20 min to denature proteins.

Reduced lysates or pre-stained protein ladder (Thermo Scientific™ #26619) were loaded into the specified lanes of a Bolt 4-12% Bis-Tris Plus gel (Invitrogen™ #NW04125BOX) in Bolt™ MES SDS Running Buffer (Invitrogen™ #B0002). A 180 V current was applied for 60 min to separate proteins by electrophoretic mobility (i.e. size). Separated proteins were transferred onto an iBlot™ 2 Transfer Stacks PVFD membrane (Invitrogen™ IB24002) using dry transfer (as described previously [53]) in an iBlot™ 2 system.

Following transfer, the PVFD membranes were incubated at room temperature on a shaker for 1 h or overnight at 4 °C in 5% BSA-supplemented TBS/0.1% Tween-20 (TBST) to block non-specific protein binding during subsequent antibody staining. Blocked membranes were then incubated at 4 °C overnight in primary antibody solution. Blots were then washed 3 x 5 min each in TBST and then incubated at room temperature for 1 hour in secondary antibody solution.

Labeled blots were washed 4 x 5 min each then incubated in SuperSignal™ West Pico PLUS Chemiluminescent Substrate (Thermo Scientific™ #34577) at room temperature for 3 min. Developed blots were scanned for chemiluminescence on a BioRad ChemiDoc Imaging System. ImageJ was used to enhance raw images for display and to quantify band intensity. Band intensity is reported as chemiluminescence of the protein of interest, normalized to the GAPDH loading control.

### Prediction of Kinases

HepG2 cells were pre-treated with the KiR inhibitor panel in Opti-MEM (Supplementary Table 1) at 500 nM for 1 h, after which media was removed, and cells were replenished with either DSM or DENV2 MON601 at an MOI of 2 in the presence of the inhibitor panel. After 90 min, cells were washed once with 1X PBS then replenished with inhibitor-supplemented DSM. 24 h post-infection, cells were fixed and stained with Env-488. The percentage of cells staining positive for Env was quantified by flow cytometry. Inhibitor-induced background fluorescence was measured on uninfected, inhibitor-treated samples and subtracted from infection values. Resulting values below zero were adjusted to zero. Grubb’s Test was used to remove outlier data.

The elastic net regularization algorithm used for this study was published previously [50; 54]. Briefly, normalized percent infection data from five independent kinase inhibitor screens and existing biochemical data of the kinase inhibitors against 300 recombinant protein kinases [47], were input into the elastic net regularization algorithm using a condition-specific cross-validation strategy. The glmnet package (version 2.2.1, https://github.com/bbalasub1/glmnet_python) in Python (version 3.7.6, https://www.python.org) was used for performing elastic net regression, with the elastic net mixing parameter α, which confers the stringency of model selection, scanned from 0.1 to 1.0 in steps of 0.1. Predictions using an α of 0.8 are reported in this manuscript. The regularization path was computed for the elastic net penalty at a grid of values for the regularization parameter λ of 10^3^. The Python code used for this analysis is provided in Supplementary Files (Supplementary File 1).

### Building Kinase Interaction Networks

Phosphosignaling networks were built to identify upstream/downstream kinases of the KiR predicted kinases and infer the connections between them. Searches for two layers upstream/downstream of the KiR predicted kinases were done using the kinase-substrate phosphorylation database PhosphoSitePlus^®^ [55]. These data were visualized using Cytoscape 3.9.1 [56]. L0, L1, and L2 nodes were manually spatially organized, then the degree-sorted circular layout algorithm was applied to spatially organize predicted RTKs and their interactions based on degree of connection.

### Statistical Analyses

Sample set details and method of statistical analysis are reported for each experiment in the corresponding figure legend. For multiple comparison analyses for data with normal distribution, One-Way ANOVA was used, while data with skewed distribution were analyzed by Brown-Forsythe.

## RESULTS

### Kinase Regression predicts multiple receptor tyrosine kinases that regulate DENV infection of hepatocytes

The liver is directly infected by DENV and can be critically damaged in severe dengue cases (reviewed in [57]). Additionally, kinase regulation of dengue-induced liver injury has been reported [40; 58]. Considering this, we chose to apply kinase regression (KiR) to DENV infection of HepG2 hepatoma cells, a widely used model for studying DENV infection of the liver. We pre-treated HepG2 cells with a panel of 38 inhibitors collectively targeting 300 kinases (Supplementary Table 1) with known target overlap at 500 nM. An hour after treatment, we infected the cells with DENV2 MON601 at an MOI of 2 and continued inhibitor treatment. Twenty-four hours post-infection, we fixed the cells and stained them with a 488-conjugated Env antibody solution. The percentage of cells staining positive for Env was quantified by flow cytometry (Supplementary Figure 1). Several kinase inhibitors dramatically decreased DENV infection while other inhibitors had little effect; in contrast, no inhibitors dramatically elevated levels of infection (Figure 1B). To predict kinases important for DENV infection, we input these data, along with existing information on kinase-substrate inhibition, into an elastic net regularization algorithm (see Methods) [47; 48]. This analysis led us to identify 36 kinases as potentially crucial in regulating dengue infection (Figure 1C, Supplementary Table 2).

To prioritize predictions for subsequent investigation, we explored which of the predicted kinases are targets of FDA approved drugs. Strikingly, the majority of predicted kinases with available drugs were receptor tyrosine kinases (RTKs), despite the fact that RTKs represent only about 10% of all kinases (Figure 1C) [59]. The predicted RTKs included Ephrin type-A receptor 4 (EPHA4), Ephrin type-B receptor 3 (EPHB3), Ephrin type-B receptor 4 (EPHB4), Receptor tyrosine-protein kinase erbB-2 (ERBB2), Fibroblast growth factor receptor 2 (FGFR2), Insulin-like growth factor 1 receptor (IGF1R), and Proto-oncogene tyrosine-protein kinase receptor Ret (RET).

To further assess the potential of RTKs as regulators of DENV infection, we examined if the seven predicted RTKs interacted with other KiR predictions or with kinases previously shown to be important for DENV infection [32; 60; 61; 62; 63; 64; 65; 66; 67; 68; 69; 70; 71; 72; 73; 74; 75; 76; 77; 78; 79; 80; 81; 82; 83; 84; 85; 86; 87; 88; 89; 90; 91]. We utilized the kinase-substrate phosphorylation database PhosphoSitePlus^®^ [55] to identify interacting kinases and substrates of the predicted RTKs and then visualized these interactions using Cytoscape 3.9.1 (Fig 1D). Known interactions for EPHA4 and EPHB4 are entirely composed of autophosphorylation events within the PhosphoSitePlus^®^ database, so they are not included in this network. Notably, the PhosphoSitePlus^®^ database does not exhaustively catalog all kinase-substrate interactions, so additional interactions beyond those illustrated in Figure 1D likely exist. Nevertheless, we observed that EPHB3, ERBB2, FGFR2, IGF1R, and RET are upstream regulators both of kinases shown to be important in previous dengue research and of kinases predicted by KiR. We therefore hypothesized that the KiR-predicted RTKs are key mediators of DENV infection.

### Elevated levels of RTKs are observed in DENV-infected cells

RTKs are stationed on the plasma membrane of cells, where they mediate growth factor signaling and cell-cell communication (reviewed in [92]). After interacting with ligand, RTKs are often endocytosed and activate signaling cascades that orchestrate cell function. There is extensive evidence that viruses can alter canonical RTK expression to hijack endocytosis or manipulate replication and cell death machinery, as reviewed in [93]. We thus investigated whether the KiR predicted RTKs – EPHA4, EPHB3, EPHB4, ERBB2, FGFR2, IGF1R, and RET – were present at higher levels in infected cells.

We infected HepG2 cells with DENV2 MON601 for 24 h and then stained them with Phycoerythrin (PE)-or Pacific Blue-labeled RTK antibodies in parallel with Env-AlexaFluor 488 and NS3-AlexaFluor 647; RTK-unstained samples were included as a control (Figure 2A). The mean fluorescence intensity (MFI) of each labeled RTK was first analyzed in uninfected cells to establish the baseline expression of each RTK. We then quantified the MFI of the Env-(bystander) and the Env+ (infected) cell populations and normalized these data to the MFI of uninfected cells for each RTK (Figure 2B). The same analysis was performed on NS3+ infected cells and NS3-bystander cells (Figure 2C). Interestingly, we found that the level of each RTK was significantly higher in infected cells compared to bystander cells. These results were consistent across biological replicates with varied infection rates (Supplementary Figure 2).

**Figure 2.**
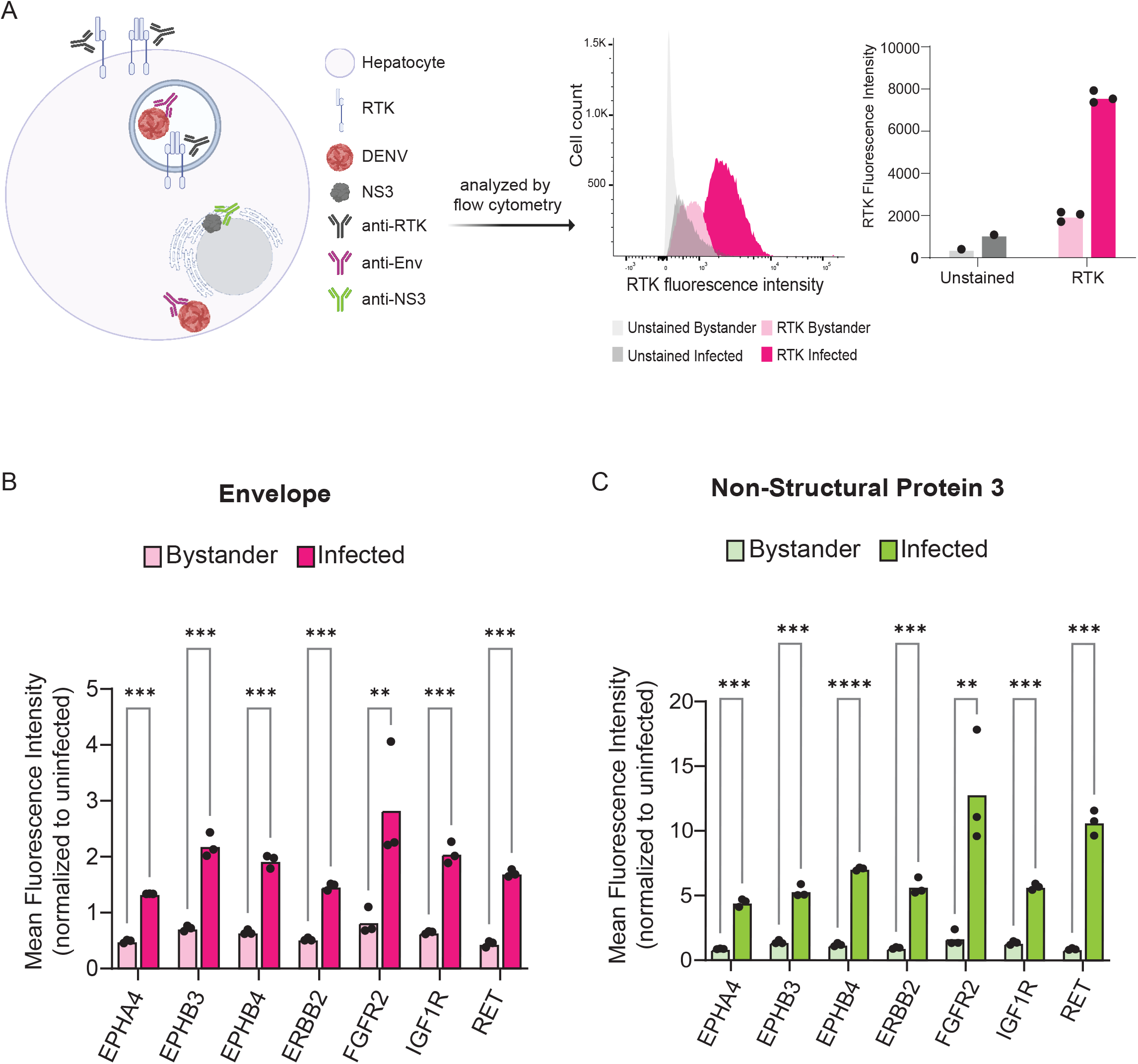
Infected cells exhibit increased levels of KiR-predicted RTKs. HepG2 cells were infected with DENV at an MOI of 2 for 24 h then analyzed for protein levels of predicted RTKs by flow cytometry. (A) Diagram of detection method for RTK expression. HepG2 cells were permeabilized to simultaneously probe for total protein levels of RTK and DENV Env or NS3. Plots on the right show a representative histogram and quantification for RTK expression analysis employed: fluorescence intensity of PE-tagged RET is shown in Env-(bystander) versus Env+ (infected) cells. Cells unstained for RTK are included as a staining control. (B) Difference in mean fluorescence intensity (MFI) of each RTK between Env-and Env+ cells or (C) NS3-(bystander) and NS3+ (infected) cells. Significance of the differences are indicated by Student’s t-test as denoted by asterisks, where ** = p-value<0.005 and *** = p-value<0.0005.

### Surface levels of RTKs are also elevated in HepG2 cells infected with DENV

RTKs can regulate infection either on the surface as a mediator of viral endocytosis or within the cell as a signal transducer. This led us to investigated whether RTK expression specifically at the cell surface was altered in infected cells. We probed 24 hr DENV-infected HepG2 cells for RTK prior to permeabilization such that only surface exposed RTK would be measured (Figure 3A). We compared the RTK MFI of bystander and infected cells for DENV Env (Figure 3B) and NS3 (Figure 3C). We found that, similar to what was observed for total RTK levels, the amount of RTK specifically at the surface was significantly increased in infected cells compared to bystander cells.

**Figure 3.**
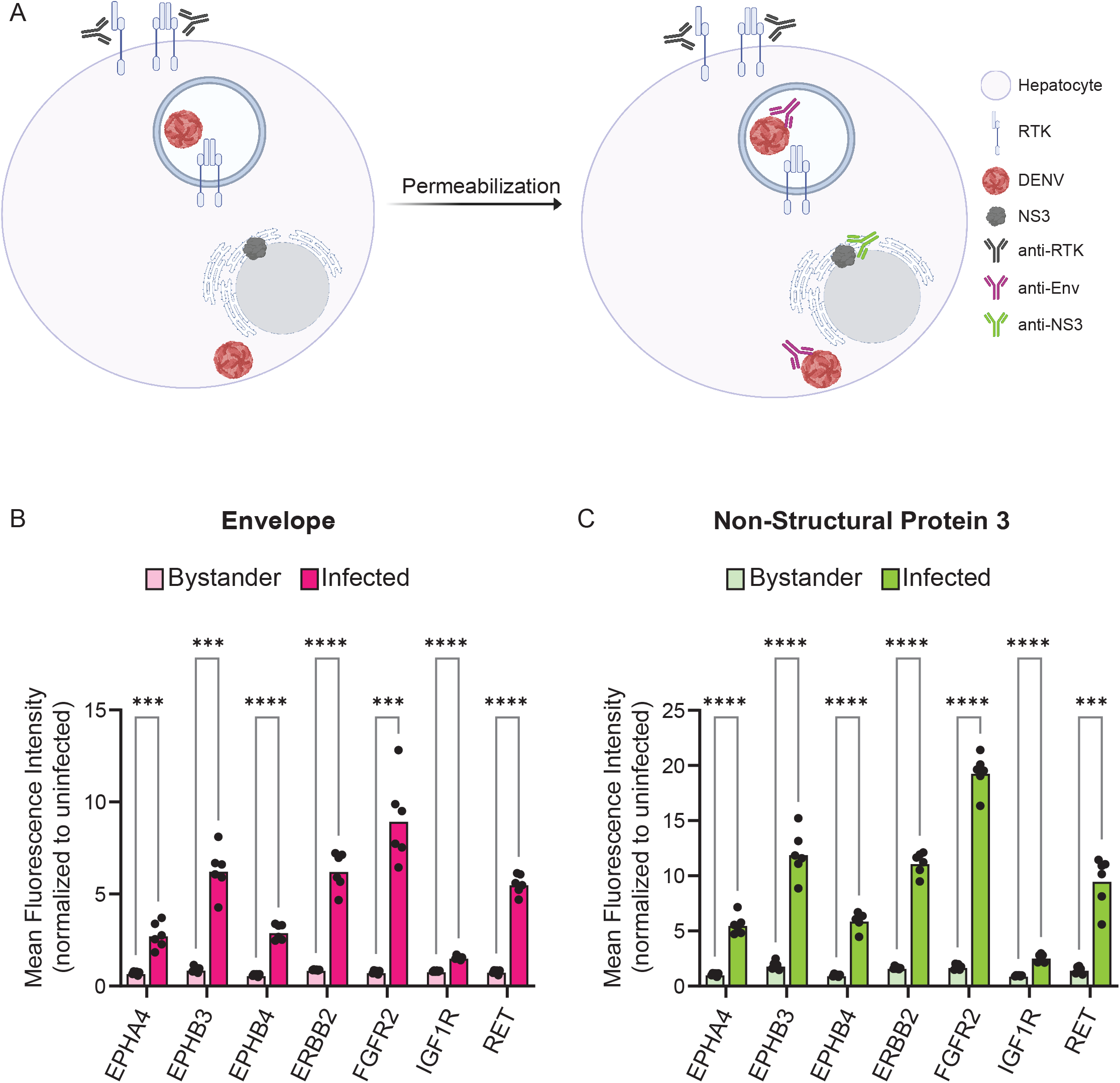
Infected cells have increased surface expression of predicted RTKs. HepG2 cells were infected with DENV at a MOI of 2 for 24 h then analyzed for protein levels of predicted RTKs by flow cytometry. (A) Diagram of detection method for RTK expression. HepG2 cells were probed for surface protein levels of RTK then permeabilized to probe for DENV Env or NS3. (B) Difference in MFI of each RTK between Env-(bystander) and Env+ (infected) cells or (C) NS3-(bystander) and NS3+ (infected) cells. Significance of the differences are indicated by Student’s t-test as denoted by asterisks, where *** = p-value<0.0005 and **** = p-value<0.00005.

### Knockdown of ERBB2, FGFR2, or IGF1R Impairs DENV Infection

Higher levels of RTK does not necessarily translate to a functional role, so we next determined if genetically reducing RTK levels affected DENV infection. We generated individual lentivirus clones carrying a puromycin selection marker and shRNA against RTK or scrambled shRNA, as a control (Supplementary Table 3). In total, we generated a single scrambled control clone and at least one clone for each RTK. We transduced HepG2 cells with each lentivirus clone and selected cells with puromycin for seven days. Cells with <10% loss in viability were analyzed by western blot to confirm protein knockdown. We successfully obtained one or more knockdown lines for EPHB4, ERBB2, FGFR2, and IGF1R (Figure 4A-B), but we were unable to procure knockdown lines for EPHA4, EPHB4, or RET, due to either loss in viability or observing no reduction in protein levels after lentiviral transduction.

**Figure 4.**
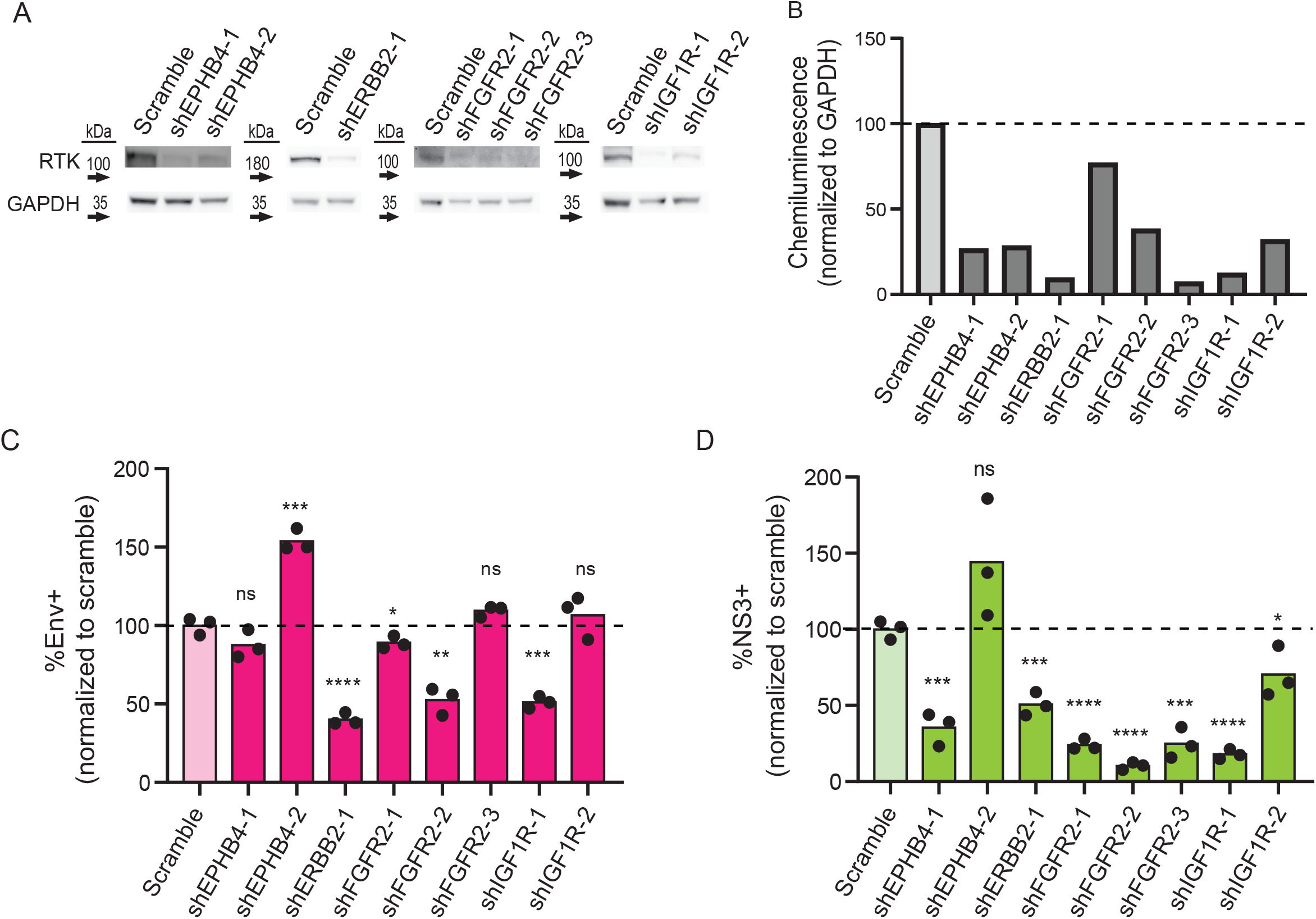
Knockdown of ERBB2, FGFR2, and IGF1R results in decreased DENV infection. HepG2 cells were transduced with shRNA lentivirus targeting KiR-predicted RTKs then analyzed for protein reduction by western blot (A-B). Viable cells with successful knockdown in protein expression were infected with DENV2 MON601, and the percent change of Env+ (C) and NS3+ (D) cells is shown. Significance of the differences are indicated by Student’s t-test as denoted by asterisks, where ns = non-significant, * = p-value<0.05, ** = p-value<0.05, *** = p-value<0.0005, and **** = p-value<0.00005.

To test whether reduced protein levels of EPHB4, ERBB2, FGFR2, or IGF1R impacted infection, we infected control and knockdown cell lines with DENV2 MON601 for 24 h. We then quantified the percentage of Env+ or NS3+ cells by flow cytometry. We calculated infection in each knockdown line as a percentage of the scramble control (Figure 4C-D). Remarkably, ERBB2 knockdown led to significantly fewer infected cells, as did reduction of IGF1R or FGFR2 in some cell lines. The effect of EPHB4 knockdown on infection varied, making its impact on infection unclear. Together, our data suggest that depletion of ERBB2, FGFR2 and IGF1R can interfere with DENV infection in hepatocytes.

### DENV infection induces ERBB2 and IGF1R phosphorylation

We next investigated if DENV infection impacts phosphorylation of the RTKs whose knockdown reduced infection. After infecting HepG2 cells with DENV2 MON601, we collected cell lysates at various time increments throughout infection. We probed cell lysates for p-ERBB2 (Thr686), p-IGF1R (Tyr1161/Tyr1165/Tyr1166), p-FGFR2 (Tyr653/654) and GAPDH; however, we did not detect a robust signal for phosphorylated FGFR2 in infected or uninfected cells (data not shown). Infection was also monitored at each time point to ensure robust infection (Supplementary Figure 4).

We observed elevated levels of phosphorylated ERBB2 and IGF1R with differing kinetic patterns during infection (Figure 5A-C). Interestingly, phosphorylated ERBB2 increased gradually from 0.5 to 1.5 hpi, having a greater than 2-fold increase at 1.5 hpi on average, then declined from 6 to 8 hpi. ERBB2 phosphorylation was increased at 16 hpi but then gradually declined through 24 hpi. In contrast, phosphorylated IGF1R was induced from 0.5 to 1.5 hpi then returned to baseline for the remainder of infection. These kinetic differences in RTK activation suggest that different RTKs have roles at distinct stages of DENV infection in hepatocytes.

**Figure 5.**
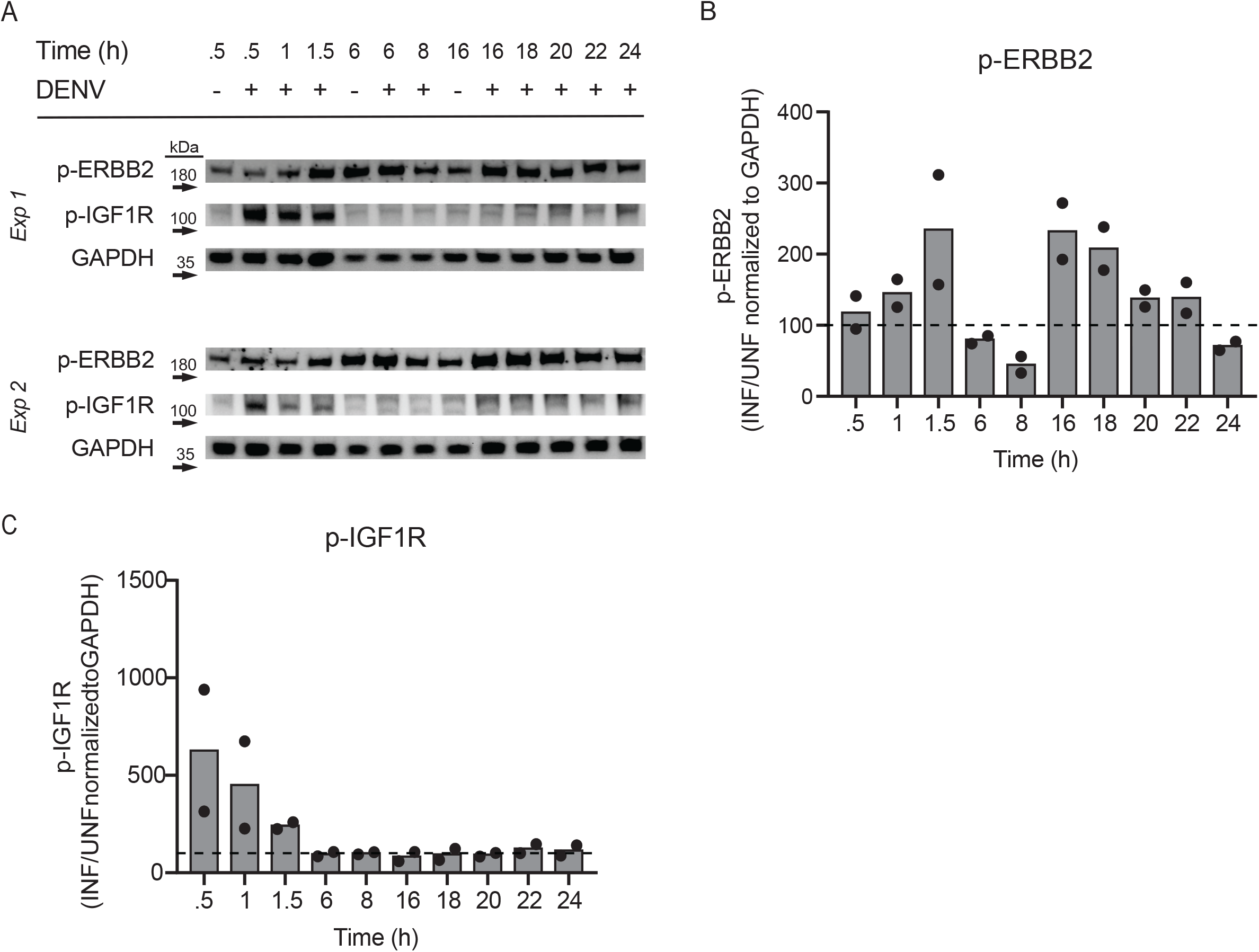
Phosphorylation of ERBB2 and IGF1R is modulated by DENV infection. HepG2 cells were infected with DENV2 MON601 then lysed at the indicated time-points. Lysates were analyzed for p-ERBB2 (A) and p-IGF1R (B) levels by western blot. p-ERBB2 and p-IGF1R band intensity was normalized to GAPDH. Percent change in normalized intensity between infected samples and appropriate uninfected control sample was graphed [0.5 – 1.5 hpi were compared to uninfected 0.5 hour sample, 6 – 8 hpi to uninfected 6 hr sample, and 16-24 hpi to uninfected 16 hr sample] (C-D). Data represent two independent experiments. Dashed line demarks the uninfected control. One-way ANOVA analysis indicated significant differences in p-ERBB2 (p=0.0155). Brown-Forsythe test indicated significant differences in p-IGF1R (p<0.0001).

## DISCUSSION

Dengue poses a significant global health threat. To mitigate this, we sought to identify host regulators of DENV infection to inform therapeutic interventions against DENV infection. We chose to focus on kinases in our study, as there is an extensive library of pharmacological tools to modulate their activity in the clinic and thus could provide useful therapeutic targets for dengue. Using kinase regression (KiR), an approach consisting of a small inhibitor screen and linear regression modeling, we predicted 36 kinases that could regulate DENV infection in HepG2 cells (Figure 1A-C). These 36 kinases, likely significant in various cell types due to the universality of kinase networks, represent potential targets for dengue interventions.

We narrowed our focus to a subset of the predicted kinases, the receptor tyrosine kinases (RTKs) – EPHA4, EPHB3, EPHB4, ERBB2, FGFR2, IGF1R, and RET – since many of these are targets of drugs already used in the clinic (Figure 1C). Interestingly, we discovered that DENV-infected HepG2 cells have elevated expression of both total and surface RTK (Figures 2-3). A limitation of this approach is that there is that flow cytometric fluorescence measurements are not always linear and thus we cannot quantify the absolute magnitude of the change in levels of RTK between infected and uninfected cells. Given this limitation, and the fact that abundance measures alone cannot verify a functional role in infection, we investigated the impact of RTK knockdown on DENV infection. We observed that depletion of ERBB2, FGFR2, or IGF1R protein led to reduced levels of both Env+ and NS3+ cells, suggesting that these RTKs can be targeted to interrupt DENV infection (Figure 4).

Strikingly, we observe DENV infection leads to phosphorylation of ERBB2 and IGF1R at different times during infection (Figure 5). This is particularly intriguing given that our data indicated that no single kinase knockdown or kinase inhibitor can, on its own, completely abolish DENV infection of HepG2 cells (Figure 1, Figure 4), and previous studies have demonstrated that combination kinase inhibitor treatment leads to increased efficacy against DENV [39; 94]. Therefore, additional research is needed to examine the different functional kinetics of ERBB2 and IGF1R during DENV infection. One hypothesis is that IGF1R activity is solely important early during infection, while ERBB2 has roles throughout infection. In this case, combination treatment to block both IGF1R and ERBB2 could result in more robust inhibition than targeting either one individually.

Of note, IGF1R and ERBB2 drug combinations are already being investigated in the context of cancer [95; 96]. Additionally, both IGF1R and ERBB2 interact with other host factors known to play a role in dengue infection or pathogenesis, such as phosphoinositide 3-kinase (PI3K)/RAC-alpha serine threonine-protein kinase (AKT) [32; 67; 75; 77; 97; 98; 99; 100], proto-oncogene tyrosine-protein kinase Src [36; 60; 61; 101], and mitogen-activated protein kinases (ERK) [20; 68; 84; 85; 102; 103]. Thus, the activity of ERBB2 and IGF1R during DENV infection should be further investigated to understand their utility as dual targets of dengue therapeutics.

In addition to small molecule inhibitors, RTKs can also be targeted by monoclonal antibodies in the clinic [104; 105]. Elevated surface levels of RTKs in infected cells (Figure 3) suggests that they could have roles as receptors for viral entry, in which case blocking receptor interaction would inhibit DENV infection. If further investigation demonstrates their role as entry receptors, approved monoclonal antibodies such as Teprotumumab (IGF1R) and Trastuzumab (ERBB2) could prove useful for blocking DENV infection.

Importantly, since kinase inhibitors have the propensity to inhibit multiple kinases through polypharmacology, single kinase inhibitors that block multiple kinase regulators of dengue should also be explored. For instance, FGFR inhibitor Futibatinib has the potential to block both FGFR2 activity demonstrated in the present study (Figure 4) in addition to FGFR4 activity previously shown to regulate DENV infection [31]. Promiscuous inhibitors targeting a combination of kinases identified in this study and others could enhance anti-DENV activity without generating toxicity often observed in multi-drug regimens.

While we focused on only a subset of predicted kinases here, the remaining predictions are also promising as druggable regulators of DENV infection. One important mechanism that should be explored is how downstream targets of KiR predicted kinases (Figure 1D) may mediate detrimental DENV immune responses which could be therapeutically blocked with RTK-targeting drugs. For instance, KiR predicted liver kinase B1 (LKB1) is part of the regulation pathway for alanine transaminase (ALT), a liver enzyme that is significantly elevated in severe dengue cases [106]. Inhibitor of nuclear factor kappa-B kinase subunit alpha (IKK-α) and NF-kappa-B-inducing kinase (NIK) are upstream regulators of TNFα which is implicated in DENV pathogenesis [107]. Additionally, rho associated coiled-coil containing protein kinase 1 (ROCK1), which is a target of Fasudil – used in the clinic for cerebral vasospasm – is known to regulate the migration and adhesion of inflammatory cells [108]. These examples highlight pathways that could be targeted to block infection while simultaneously preventing immune-mediated disease. The network analysis we utilize provides a framework for forming such hypotheses on the systematic mechanism of kinase involvement.

In conclusion, this study provides novel insight into kinase regulators of DENV infection and highlights the potential of receptor tyrosine kinases as therapeutic targets against dengue.

## ACKNOWLEDGEMENTS

This work was supported by the T32 Institutional Training Grant 5T32AI007509-20, awarded to NMB and by Seattle Children’s Research Institute internal funding. Diagrams used in this work were adapted from BioRender.com (2023). Retrieved from https://app.biorender.com/biorender-templates.

## Notes

### Competing Interest Statement

The authors have declared no competing interest.

