## Supplementary Information for "Multiple Receptor Tyrosine Kinases Regulate Dengue Infection of Hepatocytes"

**Supplementary Figure 1 Flow gating strategy for DENV infection in HepG2 cells**

DENV infection is quantified by gating for cells by size (FSC-A x SSC-A) then measuring the percent of cells staining positive for DENV envelope (Env) protein or non-structural protein 3 (NS3).

**Supplementary Figure 2 Difference in total RTK expression in DENV-infected cells across biological replicates with varying infection rates**

The fold change in total RTK expression between bystander and infected cells is shown against the average infection rate for each experiment for Env (A) and NS3 (B). Dashed line denotes fold change = 1 to draw attention to the difference in scale for fold change across the RTKs.

**Supplementary Figure 3 Difference in surface RTK expression in DENV-infected cells across biological replicates with varying infection rates**

The fold change in surface RTK expression between bystander and infected cells is shown against the average infection rate for each experiment for Env (A) and NS3 (B). Dashed line denotes fold change = 1 to draw attention to the difference in scale for fold change across the RTKs.

**Supplementary Figure 4 Time Course Infections**

HepG2 cells were infected with DENV2 then harvested at the indicated time-point to quantify infection by flow cytometry (in parallel with p-ERBB2 and -IGF1R analysis [Figure 5]). Cells were fixed, permeabilized, blocked then stained with anti-Env-488. Percent of Env+ cells for two independent experiments are shown (A-B).

Supplementary Tables

**Supplementary Table 1** lists the kinase inhibitors used in the KiR screen with corresponding CAS number.

| **Supplementary Table 1 Kinase Regression Inhibitor Panel** | |
|---|---|
| *Inhibitor ID* | *CAS #* |
| Aminopurvanolol A | 220792-57-4 |
| AMPK Inhibitor; Compound C (Dorsomorphin) | 866405-64-3 |
| Bosutinib | 380843-75-4 |
| Casein kinase I inhibitor D4476 | 301836-43-1 |
| Cdk1/2 Inhibitor III | 443798-55-8 |
| CDK2 inhibitor IV; NU6140 | 444723-13-1 |
| CDK4 inhibitor | 546102-60-7 |
| Dasatinib | 302962-49-8 |
| Dovitinib | 405169-16-6 |
| EGFR/ErbB2/ErbB4 inhibitor | 881001-19-0 |
| Erlotinib | 183321-74-6 |

| | |
|---|---|
| Gefitinib | 184475-35-2 |
| Go 6976 | 136194-77-9 |
| Go 6983 | 133053-19-7 |
| GSK inhibitor IX (BIO) | 667463-62-9 |
| GSK-3 Inhibitor X | 740841-15-0 |
| GSK-3 Inhibitor XIII | 404828-08-6 |
| GSK-3b inhibitor I (TDZD-6) | 327036-89-5 |
| H89 | 130964-39-5 |
| Imatinib | 152459-95-5 |
| JAK inhibitor I | 457081-03-7 |
| JNK inhibitor II (SP600125) | 129-56-6 |
| K252a | 99533-80-9 |
| Lapatinib | 388082-78-8 |
| Lck inhibitor | 213743-31-8 |
| Masitinib | 790299-79-5 |
| Nilotinib | 641571-10-0 |
| PKR inhibitor | 608512-97-6 |
| ROCK inhibitor (Y-27632) | 129830-38-2 |
| SB218078 | 135897-06-2 |
| Sorafenib | 284461-73-0 |
| Staurosporine | 62996-74-1 |
| Staurosporine n benzoyl | 120685-11-2 |
| SU11274 | 658084-23-2 |

| | |
|---|---|
| SU6656 | 330161-87-0 |
| Tofacitinib | 477600-75-2 |
| TWS119 | 601514-19-6 |
| Vandetanib | 443913-73-3 |

**Supplementary Table 2** lists the kinases predicted by KiR on dengue infection in hepatocytes with the corresponding coefficient of correlation at alpha = 0.8. Positive coefficient of correlation indicates the kinase is predicted to promote DENV infection, negative coefficient of correlation indicates the kinase is predicted to restrict DENV infection.

| **Supplementary Table 2 KiR-predicted Kinases Regulating DENV Infection** | |
|---|---|
| *Predicted Kinase* | *Coefficient of Correlation* |
| ACK1 | -0.04048 |
| CHK1 | -0.02247 |
| CK1g3 | -0.05048 |
| CTK_MATK | -0.00949 |
| DYRK4 | 0.294582 |
| EPHA4 | 0.020529 |
| EPHB3 | 0.020855 |
| EPHB4 | 0.009052 |
| ERBB2/HER2 | 0.076245 |
| ERK1 | -0.129 |

| | |
|---|---|
| FGFR2 | -0.0591 |
| HIPK1 | -0.06499 |
| IGF1R | 0.294032 |
| IKKa/CHUK | 0.251537 |
| JAK3 | 0.07556 |
| KHS_MAP4K5 | 0.048963 |
| LKB1 | -0.01933 |
| MAPKAPK5/PRAK | 0.171559 |
| MARK1 | 0.00079 |
| MARK4 | 0.022757 |
| NEK11 | 0.036702 |
| NEK3 | -0.03688 |
| NIK/MAP3K14 | 0.013453 |
| P38b/MAPK11 | 0.014872 |
| P38d/MAPK13 | -0.02163 |
| PAK1 | 0.178078 |
| PAK4 | -0.0126 |
| PAK5 | -0.00049 |
| PIM3 | 0.066172 |
| PKCepsilon | -0.01607 |
| PKG1a | 0.039417 |
| RET | -0.03284 |
| ROCK1 | -0.00073 |

| | |
|---|---|
| SIK2 | 0.10975 |
| SRPK1 | 0.016612 |
| TTK | -0.01278 |

**Supplementary Table 3** details the shRNA constructs used to generate kinase knockdown cells. Scrambled control was obtained from Sigma-Aldrich® (# SCH002).

| **Supplementary Table 3 MISSION shRNA Constructs** | | | | | | | |
|---|---|---|---|---|---|---|---|
| Clone ID | Oligo Seq | RefSeq ID | Gene ID | Taxon ID | Gene Description | Validated? | Validate Cell Line |
| TRCN0000010165 | CCGGTCAGTCCGTGTGTTCTATAAACTCGAGTTTATAGAACACACGGACTGATTTTT | NM_004438.3 | 2043 | 9606 | EPH receptor A4 | Yes | A549 |
| TRCN000000000 | CCGGCCCAAACCTCTTCATATTGAACTCGAGTTCAATATGAAGAGGTTTGGGTTTTT | NM_004443.3 | 2049 | 9606 | EPH receptor B3 | Yes | MCF7 |

| | | | | | | | |
|---|---|---|---|---|---|---|---|
| 6427 | | | | | | | |
| TRCN0000000006428 | CCGGGCAGTACATTGCTCCTGGAATCTCGAGATTCCAGGAGCAATGTACTGCTTTTT | NM_004443.3 | 2049 | 9606 | EPH receptor B3 | Yes | MCF7 |
| TRCN0000000001773 | CCGGCAATGGGAGAGAAGCAGAATACTCGAGTATTCTGCTTCTCTCCCATTGTTTTT | NM_004444.4 | 2050 | 9606 | EPH receptor B4 | Yes | A549 |
| TRCN0000000001774 | CCGGTGATCTGAAGTGGGTGACATTCTCGAGAATGTCACCCACTTCAGATCATTTTT | NM_004444.4 | 2050 | 9606 | EPH receptor B4 | Yes | A549 |
| TRCN000 | CCGGTGTCAGTATCCAGGCTTTGTACTCGAGTACAA | NM_001005862.1, | 2064 | 9606 | v-erb-b2 erythroblastic leukemia viral | Yes | A549 |

| | | | | | | | |
|---|---|---|---|---|---|---|---|
| 0039878 | AGCCTGGATACTGACATTTTTG | NM_004448.2 | | | oncogene homolog 2, neuro/glioblastoma derived oncogene homolog (avian) | | |
| TRCN0000039881 | CCGGCAGTGCCAATATCCAGGAGTTCTCGAGAACTCCTGGATATTGGCACTGTTTTTG | NM_001005862.1, NM_004448.2 | 2064 | 9606 | v-erb-b2 erythroblastic leukemia viral oncogene homolog 2, neuro/glioblastoma derived oncogene homolog (avian) | Yes | A549 |
| TRCN0000000366 | CCGGGCACACACTTACAGAGCACAACTCGAGTTGTGCTCTGTAAGTGTGTGCTTTTT | NM_000141.4,NM_001144914.1,NM_001144915.1,NM_001144916.1,NM_001144917.1,NM_ | 2263 | 9606 | fibroblast growth factor receptor 2 | | |

| | | | | | |
|---|---|---|---|---|---|
| | | 0011449<br>18.1,NM_<br>022970.3 | | | |
| TR<br>CN<br>000<br>000<br>036<br>7 | CCGGGCCACCAA<br>CCAAATACCAAA<br>TCTCGAGATTTG<br>GTATTTGGTTGG<br>TGGCTTTTT | NM_0001<br>41.4,NM_<br>0011449<br>13.1,NM_<br>0011449<br>14.1,NM_<br>0011449<br>17.1,NM_<br>022970.3 | 226<br>3 | 960<br>6 | fibroblast growth<br>factor receptor 2 |
| TR<br>CN<br>000<br>000<br>036<br>8 | CCGGCCGAATGA<br>AGAACACGACCA<br>ACTCGAGTTGGT<br>CGTGTTCTTCATT<br>CGGTTTTT | NM_0001<br>41.4,NM_<br>0011449<br>13.1,NM_<br>0011449<br>14.1,NM_<br>0011449<br>15.1,NM_<br>0011449<br>16.1,NM_<br>0011449 | 226<br>3 | 960<br>6 | fibroblast growth<br>factor receptor 2 |

| | | | | | | | |
|---|---|---|---|---|---|---|---|
| | | 18.1,NM_0011449 19.1,NM_022970.3 | | | | | |
| TRCN0000000424 | CCGGGCTGATGTGTACGTTCCTGATCTCGAGATCAGGAACGTACACATCAGCTTTTT | NM_000875.3 | 3480 | 9606 | insulin-like growth factor 1 receptor | Yes | A549 |
| TRCN0000000425 | CCGGCCTTGGACGTTCTTTCAGCATCTCGAGATGCTGAAAGAACGTCCAAGGTTTTT | NM_000875.3 | 3480 | 9606 | insulin-like growth factor 1 receptor | Yes | A549 |
| TRCN0000000404 | CCGGCCGCTGGTGGACTGTAATAATCTCGAGATTATTACAGTCCACCAGCGGTTTTT | NM_020630.4,NM_020975.4 | 5979 | 9606 | ret proto-oncogene | Yes | MCF7 |

| | | | | | | | |
|---|---|---|---|---|---|---|---|
| TRCN0000000405 | CCGGGCTGCATGAGAACAACTGGATCTCGAGATCCAGTTGTTCTCATGCAGCTTTTT | NM_020630.4,NM_020975.4 | 5979 | 9606 | ret proto-oncogene | Yes | MCF7 |

Supplementary Files

**Supplementary File 1** (KinaseRegression.ipynb) includes the code used to run KiR on DENV infection.

**Supplementary File 2** (NMB064_069_070_074_075_OutliersRemoved_KinasePrediction_alpharange.txt) includes the KiR output for DENV infection across the range of alphas.

**Supplementary File 3** includes the L2 phosphosignaling network from KiR.
