## Supplementary figures and images for "Multiple Receptor Tyrosine Kinases Regulate Dengue Infection of Hepatocytes"

### Supplementary Figure 1

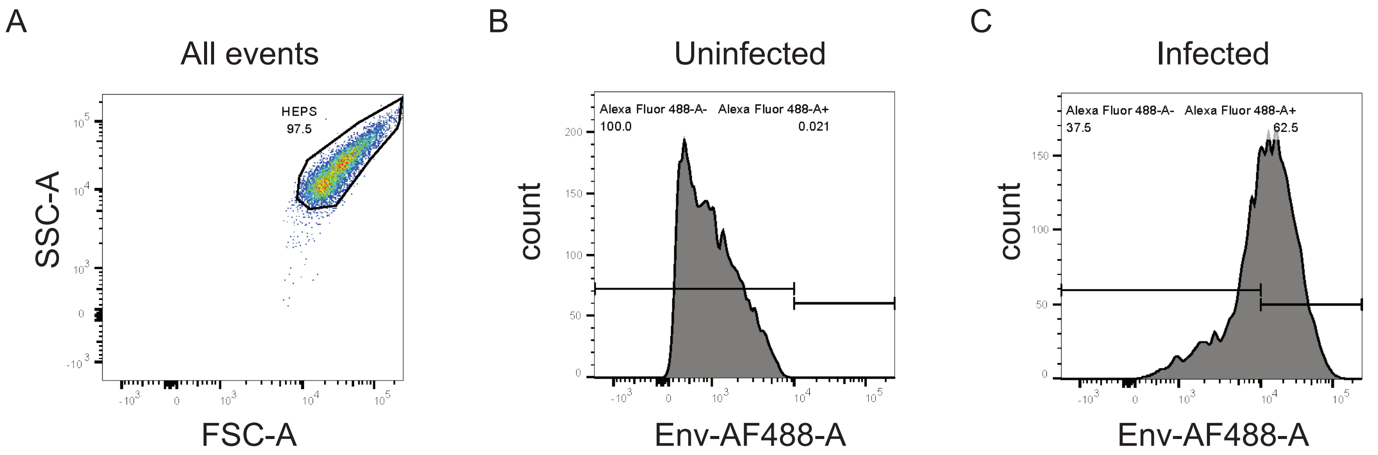

### Supplementary Figure 2

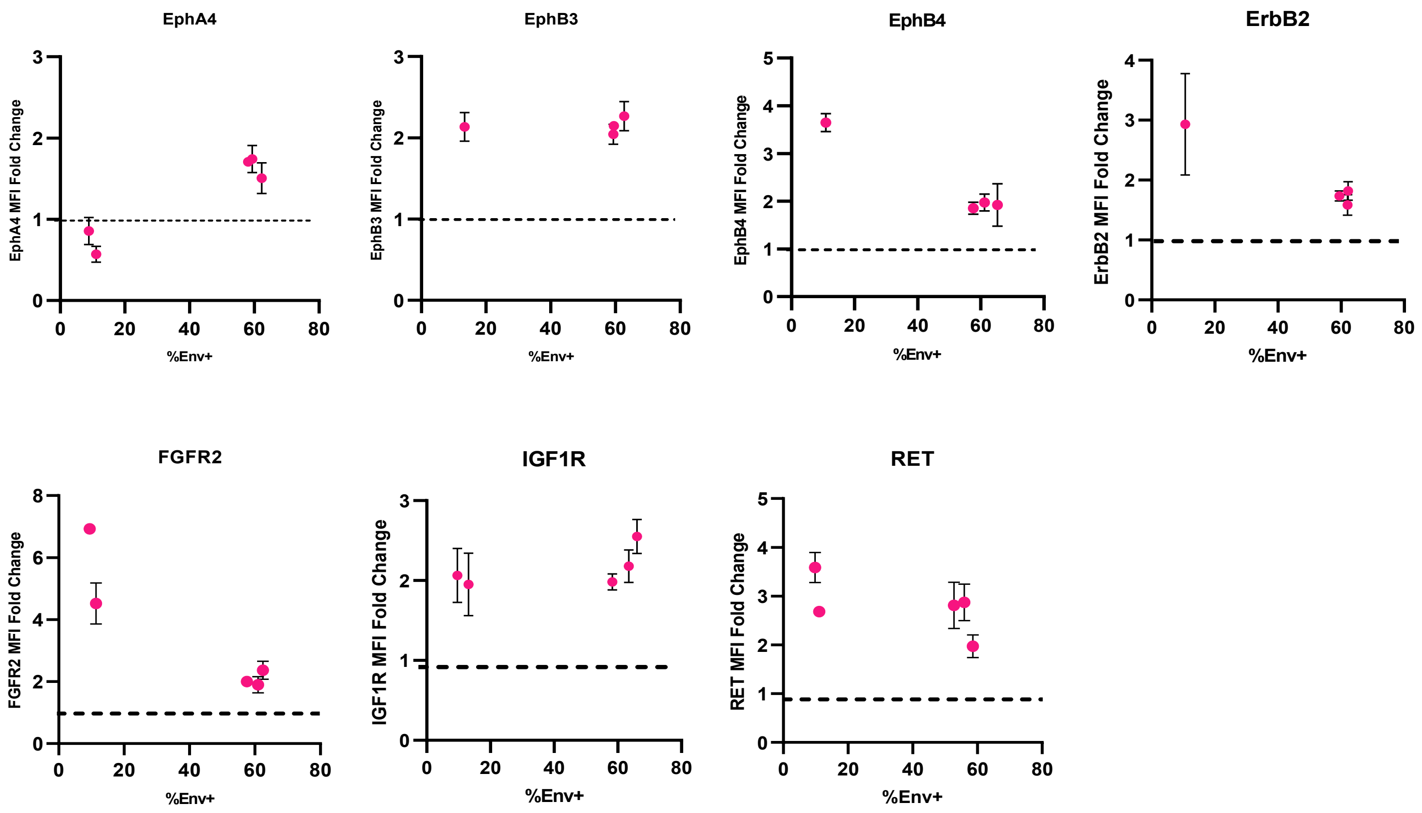

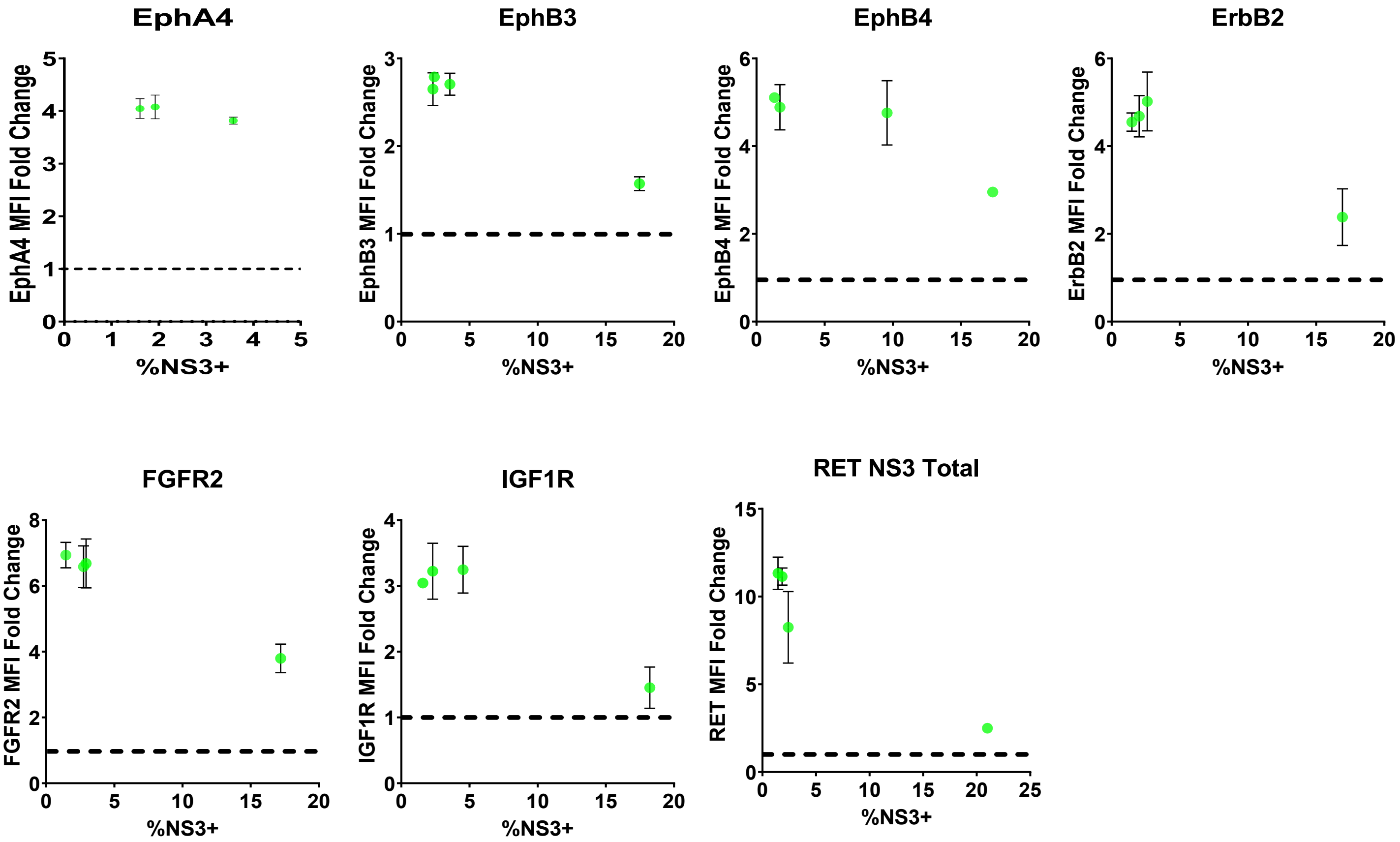

### Supplementary Figure 3

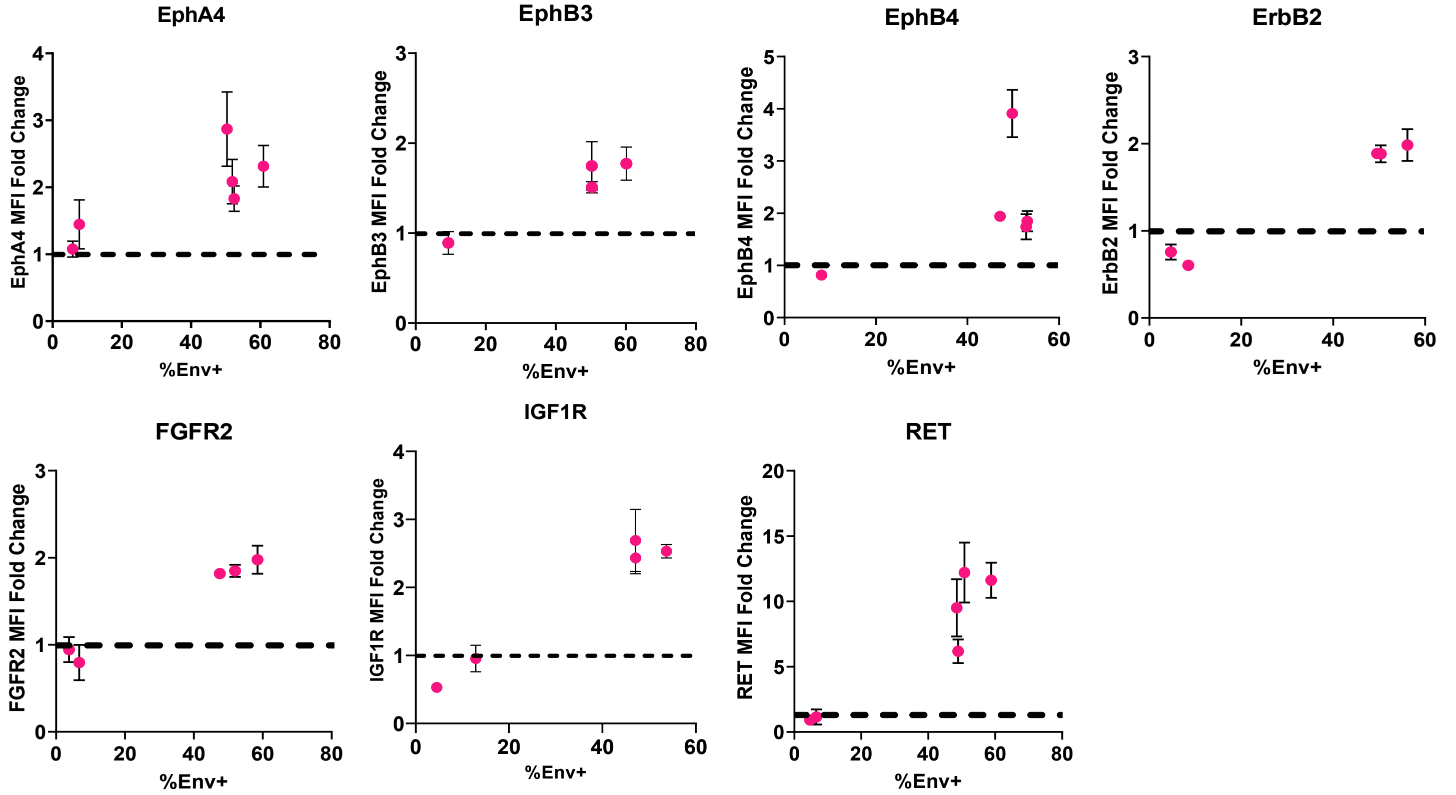

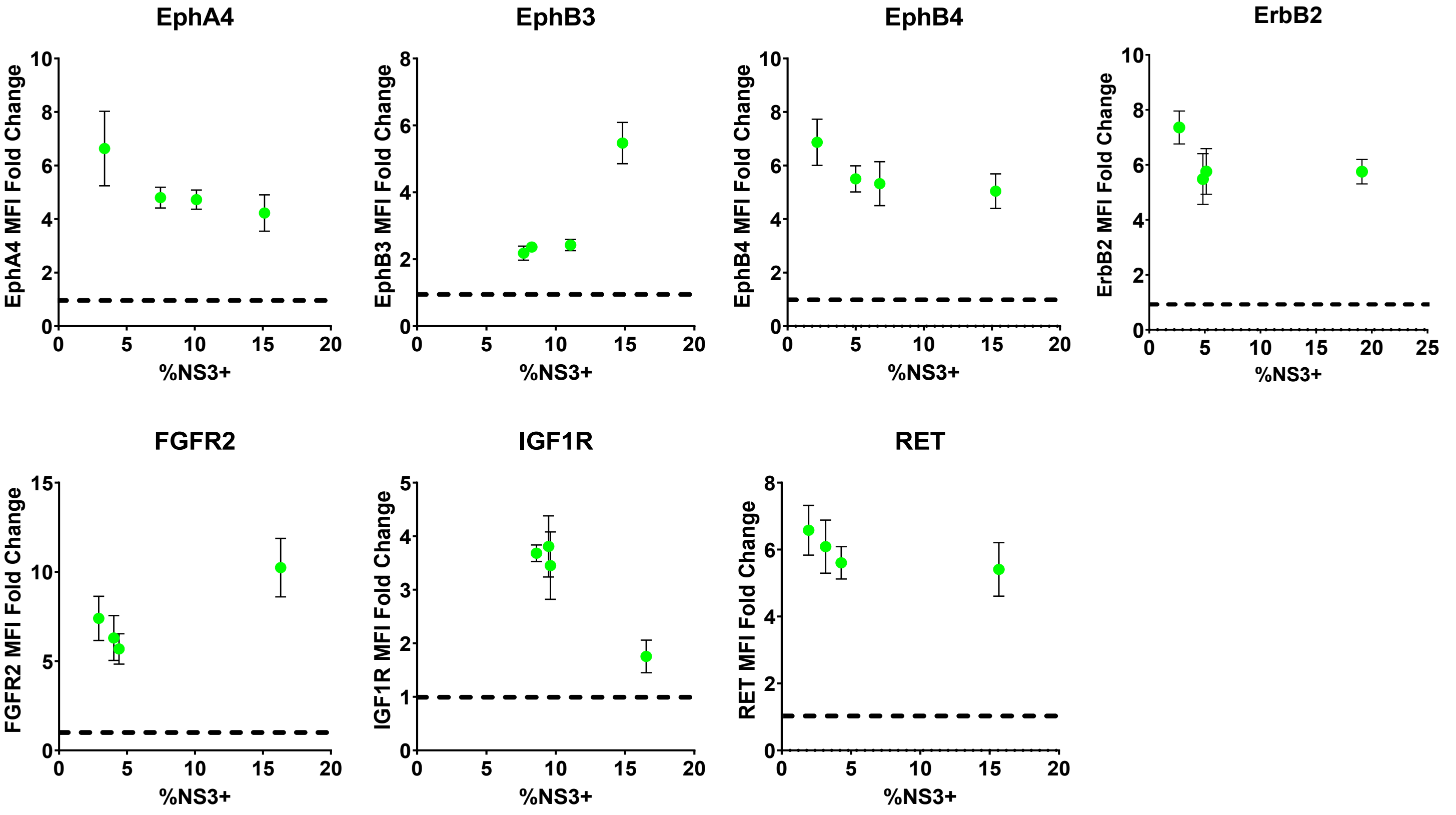

### Supplementary Figure 4

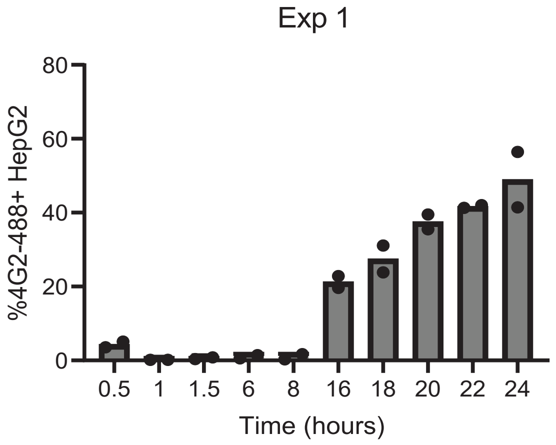

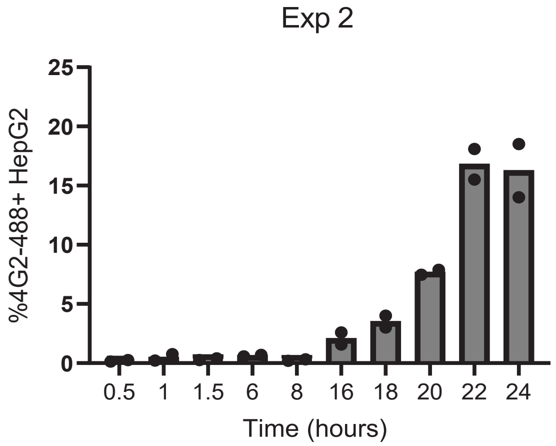
