## Supplementary material for "Multiple Receptor Tyrosine Kinases Regulate Dengue Infection of Hepatocytes": KinaseRegression.ipynb

```
# %% [code]
import sys
sys.path.append('/Users/lwei/glmnet_python')
sys.path.append('/Library/Python/2.7/site-packages')

import numpy as np
import pandas
from pandas import DataFrame
import matplotlib
import matplotlib.pyplot as plt
import matplotlib.colors as colors
from sklearn.linear_model import enet_path, ElasticNetCV
from sklearn.discriminant_analysis import LinearDiscriminantAnalysis, QuadraticDiscriminantAnalysis
from sklearn.ensemble import RandomForestClassifier
from sklearn.datasets import make_classification
from numpy import genfromtxt
import scipy
from scipy import sparse
from scipy import stats
import random
import seaborn as sns; sns.set()
from minisom import MiniSom
from matplotlib.gridspec import GridSpec
import csv
import networkx as nx
import glob

import glmnet_python
from glmnet import glmnet 
from glmnetCoef import glmnetCoef
from glmnetPredict import glmnetPredict
from glmnetPrint import glmnetPrint
from cvglmnet import cvglmnet
from cvglmnetCoef import cvglmnetCoef
from cvglmnetPredict import cvglmnetPredict
from cvglmnetPlot import cvglmnetPlot

# %% [code]
# Cross-validation plus prediction

path = "Ying/CsA/"

data = pandas.read_csv(path + "kidney.CsA.txt", sep='\t')
print(data.shape)

drugs = data['Drug']
drug_list = drugs.unique()

x = data.iloc[:, 2:].to_numpy()
y = data['Measurement'].to_numpy()

all_drugs = pandas.read_csv("Beno/178drugs_data_v2.txt", sep='\t')
print(all_drugs.shape)

x_test = all_drugs.iloc[:, 1:].to_numpy()

alpha = 0.8
nlambda = 1e3

fit = glmnet(x = x.copy(), y = y.copy(), family = 'gaussian', alpha = alpha, nlambda = nlambda)

lambdas = np.array(fit['lambdau'])

mse = np.zeros((len(lambdas), len(drug_list)))
mae = np.zeros((len(lambdas), len(drug_list)))

for index, i_lambda in enumerate(lambdas):
    i = 0
    for drug in drug_list:

        data_validate = data.drop(drugs[drugs != drug].index)

        x_validate = data_validate.iloc[:, 2:].to_numpy()
        y_true = data_validate['Measurement'].to_numpy()

        y_predict = glmnetPredict(fit, x_validate, ptype = 'response', s = scipy.float64([i_lambda]))
            
        mse[index, i] = np.sum(np.square(np.subtract(y_predict, y_true)))
        mae[index, i] = np.sum(abs(np.subtract(y_predict, y_true)))
        
        i += 1

lambda_min = lambdas[np.argmin(np.mean(mse, axis=1))]
print(np.argmin(np.mean(mse, axis=1)), lambda_min)

coef = glmnetCoef(fit, s = scipy.float64([lambda_min]), exact = False)
print('Non-zero coefficients: ', np.count_nonzero(coef[1:]))
np.savetxt("Ying/results/CsA_20200515/updated/kidney.CsA-glmnet-alpha0.8.txt", coef[1:], delimiter='\t')

prediction = glmnetPredict(fit, x_test, ptype = 'response', s = scipy.float64([lambda_min]))
np.savetxt("Ying/results/CsA_20200515/updated/kidney.CsA-glmnet-alpha0.8.prediction.txt", prediction, delimiter='\t')
```
